# Creating Aesthetically Pleasing SBGN Visualisations for Presentation and Exploration

**DOI:** 10.1101/2023.12.23.573191

**Authors:** Tobias Czauderna, Falk Schreiber

## Abstract

Visualisations are widely used to present, assess, and convey data and understanding in systems biology and other fields, and graphical representations of data and information should be precise, visually engaging, informative, and aesthetically pleasing. The Systems Biology Graphical Notation (SBGN) is a standard for visual representations of processes and networks in systems biology in the form of SBGN maps. It defines graphical elements and connection rules for knowledge representation in the form of maps. Whereas the form of the map elements is defined, those map elements have to be placed in a proper way in order to obtain useful and aesthetically pleasing visualisations. Here, we present layout considerations and guidelines for SBGN maps, as well as workflows for SBGN-ED as a tool for crafting visually pleasing SBGN visualisations for presentation and exploration.

## I. Introduction

The importance of visualisation in biology has grown significantly, as visualisations allow researchers to comprehend data and facts more effectively and to gain new perspectives on biological processes. Various visualisation techniques and overviews of methods have been discussed in the literature, including [50, 74, 129].

The task of understanding cellular or organismal processes and actions often involves the visualisation of biological networks, such as metabolic, regulatory, and signalling pathways, protein interaction networks, hormonal networks, and other complex biological interactions. Network visualisations can not only simplify complex biological data and help in their interpretation, but also foster information exchange and collaborations. Consequently, pathway databases usually contain network visualisations, and there is a variety of methods and tools to compute visual representations of biological processes.

It is essential to have a clear and accurate representation of data and information in systems biology and beyond. To do this, graphical maps must be precise, visually attractive, informative, and aesthetically pleasing. The Systems Biology Graphical Notation (SBGN) is a standard for information representation in systems biology and provides graphical elements and connection rules for knowledge representation in the form of SBGN maps. Nevertheless, it only provides some general guidelines for the arrangement of these elements, even though the correct layout is essential to ensure both practicality and beauty in these visualisations.

Here, we investigate layout methods and guidelines for SBGN maps and present workflows for SBGN-ED as a tool for building visually pleasing SBGN visualisations for both presentation as well as exploration. This paper is structured as follows: In Sect. I we introduce SBGN and COMBINE, in Sect. II we summarise important aspects of the SBGN specifications regarding visualisation rules and discuss general network layout. Section III presents tools for creating and working with SBGN maps, and in Sect. IV we discuss SBGN-ED and its usage for creating visually pleasing SBGN visualisations. We conclude with Sect. V.

### A. SBGN

The Systems Biology Graphical Notation (SBGN) [66, 86] is a standardised visual notation that represents biological systems at various levels of detail. The goal of SBGN is to provide an unambiguous graphical representation such that complex biological systems and interactions are more understandable, and to facilitate the exchange of information, knowledge, and models. SBGN has been continuously developed for more than 15 years and is widely used, for example, in databases such as BioModels [87, 92], Reactome [53, 95], Panther Pathways [98] and Pathway Commons [120], as well as in modelling software and SBGN editors such as BioUML [80], CellDesigner [43], PathVisio [85], Newt Editor [10] and SBGN-ED [26].

There are three main types of maps in SBGN:

- Process Description (PD) [123]: This map type emphasises the transformation of so-called species (molecules, complexes, perturbing agents such as external factors affecting the system, etc.). It is used to show the temporal courses of biochemical reactions and interactions in a network. Common components include: molecules (such as proteins, genes, small molecules), processes (such as chemical reactions, transport), and interaction arcs (indicating the influences molecules have on processes).
- Entity Relationship (ER) [144]: This type of map focuses on influences between the so-called entities (which are participants in relationships such as macromolecules, perturbations and phenotypes) without considering the temporal aspect. It is often used to represent regulatory or protein interaction networks, and to show the overall possible interactions of an entity. Common components include: entities and influence arcs (showing how one entity affects another, such as stimulation or inhibition).
- Activity Flow (AF) [99]: This type of map represents the influences of activities on each other, focusing on the dependencies without detailing the underlying molecular processes. It is more abstract than the PD and ER notations. Common elements include: activities (indicating some action or function of a molecule) and influence arcs (showing the effects between activities).

For each type of map, specific symbols represent different biological entities and interactions. These symbols and their usage (e. g. possible connections between them) are standardised to make it easier for people familiar with SBGN to understand the maps created by others. A typical SBGN map is shown in Fig. 1.

**Fig. 1:**
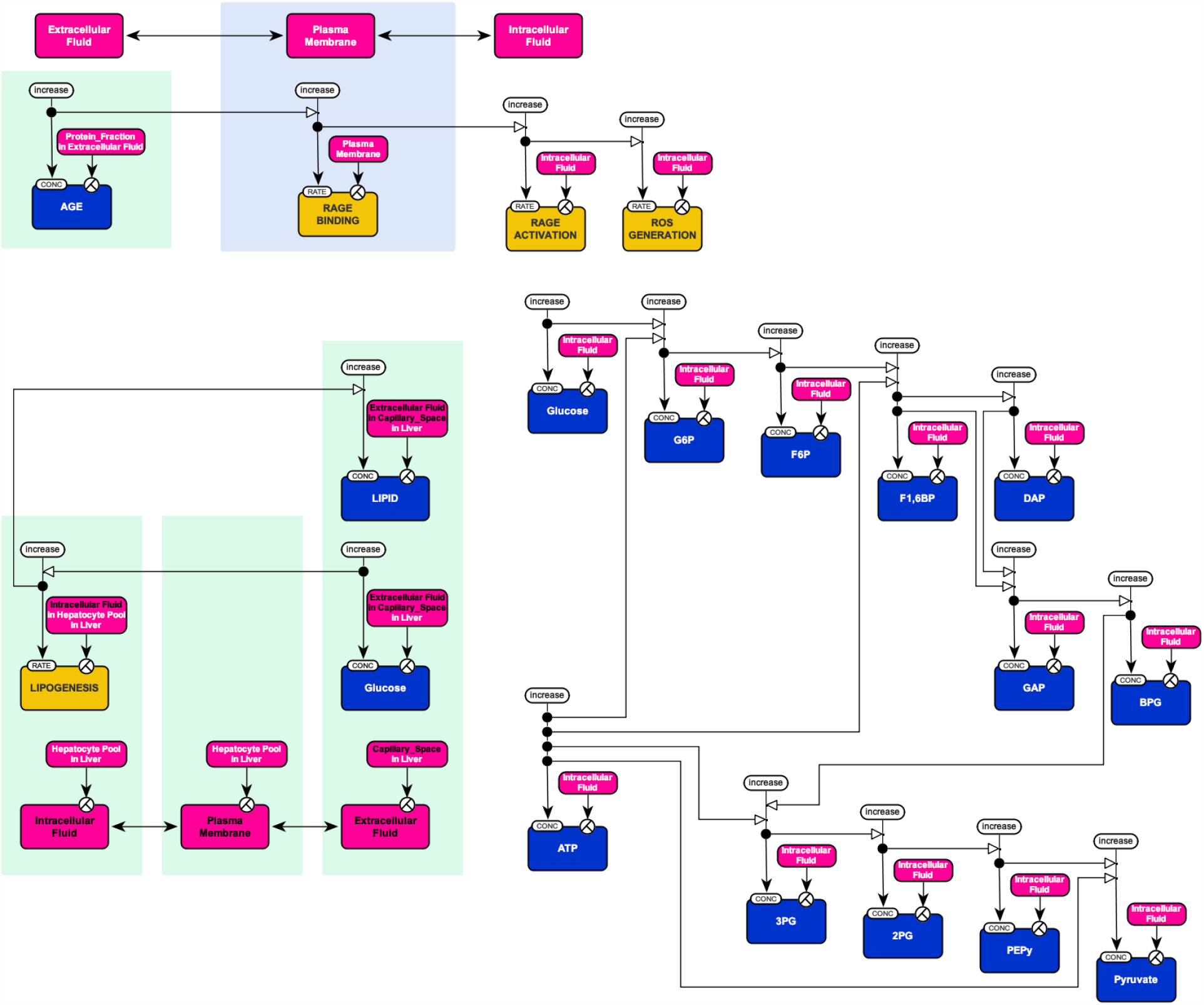
Visualisation in biology - an example using SBGN. This map outlines some of the steps in the development of diabetic retinopathy. It is a partly redrawn map of a submission of the 2010 SBGN competition (Best SBGN map: Breadth, accuracy, aesthetics) [28].

### 1) Syntax

The syntax of the three SBGN languages defines how the SBGN elements can be assembled into a valid SBGN map. It specifies which SBGN nodes (glyphs) may be connected by which SBGN arcs. This is described for each of the languages by an incidence matrix (see language specifications). For the Process Description and Activity Flow languages, it is further defined which nodes can be drawn on top of each other (containment). Moreover, there are additional syntactic rules for all three languages that are not derived from the incidence matrix. For example, in the specification for AF, it is additionally described that all elements of the language may be drawn on a Compartment node (containment) and that a Biological Activity node may have at most one Unit of Information.

### 2) Semantics

The semantic rules of the three SBGN languages describe the meaning of an SBGN map. Semantics is important so that an author can create a SBGN map that represents their understanding of a biological process or system, and a reader can understand a map without additional help. For the Activity Flow language, assuming that a biological activity has a rate, following semantic rules are defined:

- A biological activity is assigned to a single compartment, despite potentially overlapping multiple Compartment glyphs. The assignment is determined by the map’s author or the software application.
- A Positive Influence increases the rate of a biological activity.
- A Negative Influence decreases the rate of a biological activity.
- With an Unknown Influence, the effect and thus the change in rate of a biological activity is unknown.
- Only one Necessary Stimulation arc can end at a Biological Activity glyph; multiple Necessary Stimulation arcs must be linked via an AND or an OR glyph.

### B) COMBINE

The development of SBGN is part of a larger movement within systems biology to make computational and theoretical models more accessible and reproducible. COMBINE [63, 103, 154], which stands for the “COmputational Modeling in BIology” Network, is an international initiative which coordinates the development of community standards in systems biology and closely related fields. The main goal of COMBINE is to facilitate the adoption of open community standards, thus enhancing model sharing and reproducibility, which is supported by annual workshops and tutorials, and the publication of a regular special issue regarding the latest developments of COMBINE standards [130–132, 136, 138, 139]. Before COMBINE, several standardisation efforts were developed somewhat in isolation from each other. COMBINE aims to bring together these communities to ensure that standards are harmonious, non-overlapping and interoperable. Therefore, the COMBINE initiative has been instrumental in advancing systems biology by ensuring that tools, models, and data used in research can be easily shared, understood, and reproduced across the global community.

Here are the main standards and initiatives associated with COMBINE:

- SBML (Systems Biology Markup Language) [62, 73]: A format for representing biochemical reaction networks. It is a widely adopted standard in systems biology for sharing and publishing models.
- CellML [24, 25]: a format focusing on storage and exchange of computer-based mathematical models in biology, which can range from simple equations to complex cellular models.
- SBGN (Systems Biology Graphical Notation): As mentioned above, a visual notation for depicting biological processes, relationships, and activity flows.
- SED-ML (Simulation Experiment Description Markup Language) [13, 153]: A format to describe simulation experiments in a way that makes them reproducible. This means defining initial conditions, model references and outputs, among other things.
- BioPAX (Biological Pathway Exchange) [30]: A format to represent biological pathways and networks at the molecular and cellular level.
- NeuroML [54]: A format for describing models of neurons and their interconnections (networks of neurons), aimed at the neuroinformatics community.
- COMBINE Archive [12]: A single file format that can encapsulate all the different data types and formats that might be used in a computational modelling project. It makes sharing and publishing a complete model (with all associated files) more straightforward.

## II. SBGN visualisation guidelines

Important objectives for visualising all kinds of biological networks including those represented by SBGN maps include tracing the flow of information or matter through the networks, understanding how cellular processes are regulated by external or internal signals, understanding interconnections between entities such as genes, proteins, and metabolites, identifying genes that regulate larger gene sets, and finding primary and alternate paths. For the visualisation of biological systems key aspects include:

- *Pathways:* It is crucial to define the primary sequence or the flow path of substances or information, highlighting the sequence of events in their chronological order.
- *Compartments:* Biological processes occur in various compartments of the cell, and visually differentiating these areas is essential for accurate representation.
- *Complexes:* The formation of molecular complexes is a common aspect of biological processes. It is important to illustrate both the structure of these complexes and the interactions between the molecules that constitute them.

Initial guidelines are already given in the specifications of PD, ER, AF. Furthermore, in [134] design considerations for the visual representation of information from systems biology using SBGN are discussed, in particular regarding the use of SBGN glyphs.

### A. Layout guidelines

The layout guidelines describe requirements and recommendations for how SBGN maps should be presented, focusing on appearance and aesthetics. Layout requirements must be met, and layout recommendations should be met. For example, for AF maps we have the following:

#### a) Layout requirements for an AF map

- Glyphs must not overlap, with the exception of Compartment glyphs and Biological Activity glyphs overlapping Compartment glyphs.
- If an arc crosses a glyph, then the arc must be drawn on top of the glyph.
- Arcs must not overlap the border lines of a glyph.

#### Algorithm 1

Sketch of the Sugiyama (layered or hierarchical layout) algorithm for network layout (pseudocode).

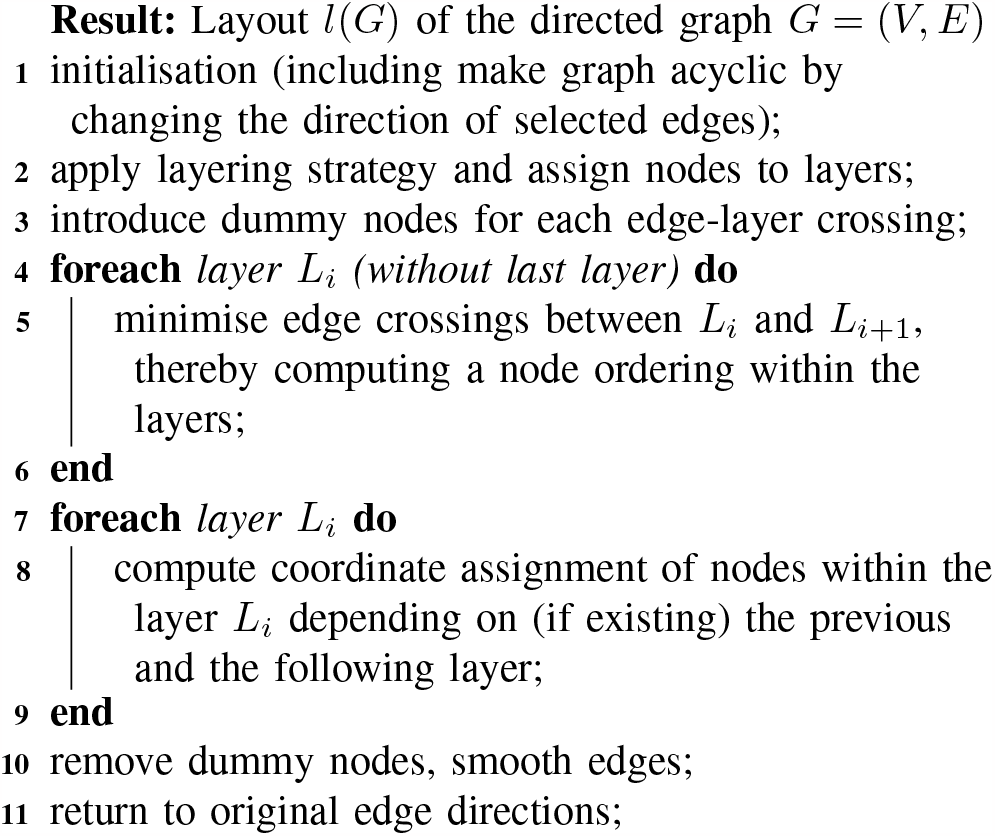
- Glyphs must be aligned horizontally or vertically.

#### b) Layout recommendations for an AF map

- The name of a glyph should be horizontally aligned and, if possible, displayed entirely within the glyph.
- Arcs should not cross glyphs.
- The number of arc crossings should be minimised.

### B. Network layout

Network layout is a critical aspect for aesthetically pleasing SBGN visualisations. It refers to the way in which the nodes (or glyphs, representing entities like proteins, genes, or metabolites) and their connections (edges or arcs, representing interactions or relationships) are arranged visually. Effective network layouts help in understanding the complex relationships and functionalities within biological systems. A layout can be produced manually, but for large networks or large amounts of networks (such as in databases) automatic layout is desired.

A network (or graph) *G* is defined as a pair of sets (*V, E*) with *V* being the set of nodes (or vertices), *V* = {*v*_1_, …, *v*_*n*_} and *E* the set of edges (or arcs), *E* = { (*v*_*i*_, *v*_*j*_) : *v*_*i*_, *v*_*j*_ ∈ *V* }. In the context of SBGN we consider edges as directed (from *v*_*i*_ to *v*_*j*_). A graph layout algorithm is a computational method used to arrange the nodes and edges of a graph in a two-dimensional (2D) or three-dimensional (3D) space by assigning coordinates to nodes and straight-lines or splines to edges. Key methods for network (or graph) layout [31, 72] include:

#### a) Layered / hierarchical layouts

This method organises nodes into distinct layers (often arranged as horizontal layers and the direction from top to bottom) [38, 46, 147]. It is particularly useful for depicting hierarchical structures, such as those seen in gene regulatory networks. In Alg. 1, a typical layered layout algorithm is sketched.

##### Algorithm 2

Sketch of the force-directed layout algorithm for network layout (pseudocode).

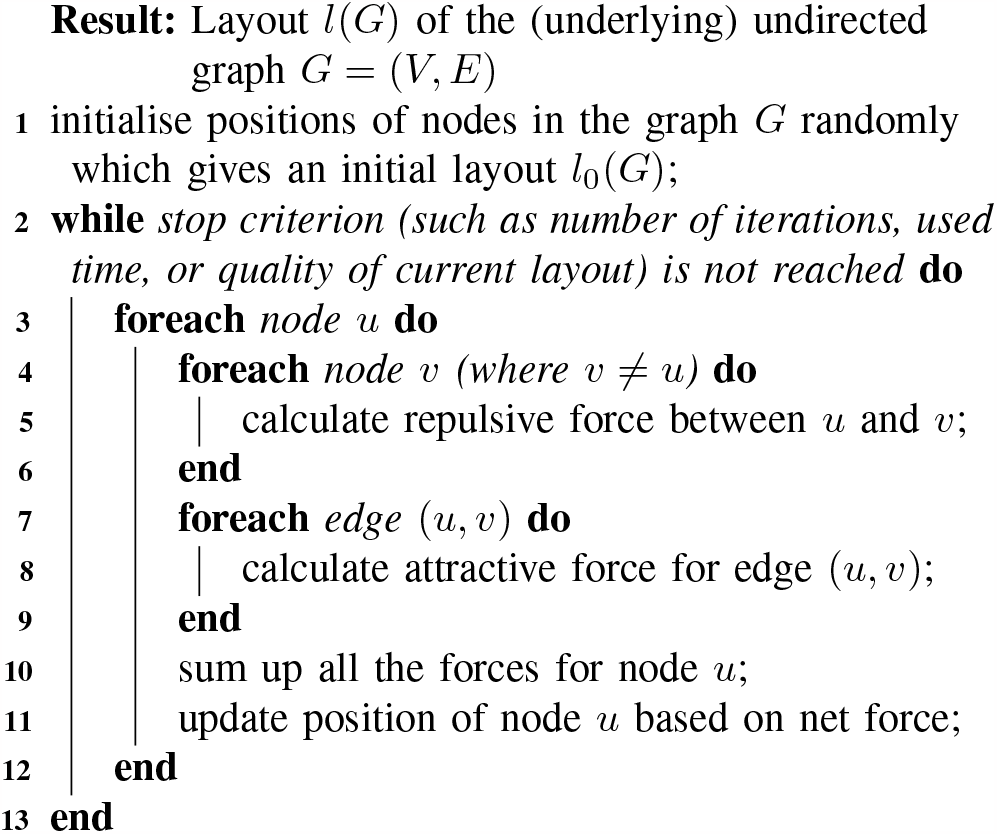

#### b) Force-directed layouts

These are layouts in which nodes repel each other like charged particles if not connected by an edge, while edges act like springs that hold the nodes together aiming for an optimal distance [37, 41, 47, 146]. This method is effective in illustrating the structure and clustering within networks, such as protein-protein interaction (PPI) networks. It is easy to implement and is probably the most often used layout method, and many variants have been proposed, in particular to speed up the computation and to include constraints into the layout (examples include [9, 39, 48, 68, 91, 106]) In Alg. 2, a typical force-directed layout algorithm is sketched.

#### c) Orthogonal layouts

This method [15, 16, 148, 157] is characterised by its use of straight lines for edges, which typically run horizontally and vertically, forming right angles. This makes the layout particularly clear and easy to follow when the graph has a complex interconnection of nodes. The nodes are placed (sometimes on grids) so that their connecting edges can be drawn as straight horizontal or vertical lines as much as possible. The challenge lies in optimising the placement of nodes and routing of edges to minimise crossings and bends.

#### d) Application domain related layouts

There are many algorithms tailored to specific biological networks, such as metabolic network layouts, e. g. [35, 71, 78, 127, 128] (where nodes represent metabolites and enzymes, while edges show metabolic reactions; often layered and force-directed methods or their combination is used to depict the flow and interconnectivity of metabolic pathways), protein interaction network layouts, e. g. [11, 40, 51, 58, 59] (where nodes represent proteins and edges their interaction; force-directed layouts are commonly used here, helping to visualise interactions and to identify key proteins that may play central roles in cellular processes), or more general overlapping and heterogeneous biological networks (multilayer networks), e. g. [45, 79, 88, 135, 137] (where networks combine different types of interactions or entities, such as networks that include both gene-gene and gene-protein interactions; hybrid layouts, combining aspects of both layered and force-directed methods, and 2.5D stacking of layouts are often employed). Overviews are also presented in [7, 75, 145].

#### e) Aesthetics and mental map

Aesthetics in graph layout (graph drawing) is crucial for clarity and engagement, emphasising principles like minimal edge crossings or showing symmetry, and several studies investigate metrics for graph layout aesthetics or deal with improving aesthetics to obtain better layouts of graphs [61, 113–115, 117]. Mental map preserving in graph layout refers to visualisation approaches that aim to maintain the user’s mental model of the graph’s structure, even when the layout of the graph is modified or updated. This concept is crucial in interactive and dynamic graph visualisation, where changes to the graph (like adding or removing nodes and edges) occur over time. Preserving the mental map helps users to understand and track these changes without losing their orientation or understanding of the overall structure of the graph [4, 49, 81, 101, 116, 118]. General open graph layout problems are also discussed in [3, 17].

## III. Tools for and usage of SBGN

Software support for SBGN and software libraries are essential for facilitating the visualisation and interpretation of complex biological networks in systems biology, enabling researchers to create, edit, and analyse detailed maps of cellular processes. A brief overview of databases using SBGN is shown in Table I including the types of SBGN maps used by the repositories and if the SBGN-ML file format [14] is supported. Software tools built in the last years are presented in Table II, including information about supported functionalities and supported SBGN map types. For a regularly updated list of tools and databases, see the SBGN web page [126]. In addition, LibSBGN [150] is a software library for reading, writing, and manipulating SBGN maps represented as files using the SBGN-ML format [14]. It is available in C++ and Java, and there is a JavaScript version [90] and a Python version [89] available.

**TABLE I:** Overview of databases and collections providing SBGN maps in any of the three languages and as SBGN-ML files (“+” supported, “-” not supported).

| Repository | PD | ER | AF | SBGN-ML |
| --- | --- | --- | --- | --- |
| AlzPathway [102] | + | - | - | - |
| AsthmaMap [96] | + | - | + | + |
| Atlas of Cancer Signalling Networks [5] | + | - | - | + |
| BioModels Database [87, 92] | + | - | - | - |
| COVID-19 Disease Map [109] | + | - | - | - |
| Metabolism Regulation Maps [97] | + | - | - | + |
| MetaCrop [56, 133] | + | - | - | + |
| NaviCell [104] | + | - | - | - |
| PANTHER Pathway [149] | + | - | - | + |
| Parkinson’s disease map [42] | + | - | - | - |
| Path2Models<sup>a</sup> [20] | + | - | - | + |
| Pathway Commons [22, 120] | + | - | - | + |
| PathWhiz [112] | + | - | - | + |
| QSDB [77] | + | - | - | - |
| Reactome [95, 100] | + | - | - | + |
| Rheumatoid Arthritis Map [140] | + | - | - | + |
| RIMAS<sup>b</sup> [64] | + | - | - | - |
| SubtiWiki [111] | + | - | - | - |
| Virtual Metabolic Human ReconMaps [105] | + | - | - | - |

<sup>a</sup>Path2Models is part of the BioModels database.
<sup>b</sup>RIMAS is no longer available, but the data is still available within Vanted.

**TABLE II:** Overview of software tools supporting 1) SBGN maps in any of the three languages, 2) translation of KEGG diagrams to SBGN maps, 3) translation between (some of) the three languages, 4) SBGN Bricks and 5) SBGN-ML (“E” editor, “V” viewer, “+” supported, “-” not supported).

| Software | V/E | PD | ER | AF | KEGG →<br>SBGN | SBGN →<br>SBGN | SBGN<br>Bricks | SBGN-ML |
| --- | --- | --- | --- | --- | --- | --- | --- | --- |
| Arcadia<sup>a</sup> [151] | V | + | - | - | - | - | - | - |
| BIOCHAM [21] | E | + | - | - | - | - | - | - |
| BioUML [80] | V | + | - | - | - | - | - | - |
| CellDesigner [94] | E | + | - | - | - | - | - | + |
| ChiBE [6] | E | + | - | - | - | - | - | - |
| COPASI [60] | V | + | - | - | - | - | - | - |
| CySBGN [55] | V | + | + | + | - | - | - | + |
| EPE<sup>a</sup> [143] | E | + | + | + | - | - | - | - |
| geneXplain platform [52] | E | + | - | - | - | - | - | - |
| JWS Online [107] | V | + | - | - | - | - | - | - |
| Krayon for SBGN [84] | E | + | - | - | - | - | + | + |
| Netbuilder’ [156] | E | + | - | - | - | - | - | - |
| Newt Editor [10] | E | + | - | + | - | - | + | + |
| Omix [34] | E | - | - | - | - | - | - | + |
| PathVisio (SBGN plugin<sup>a</sup>) [85] | E | + | + | + | - | - | - | + |
| SBGN-ED [26] | E | + | + | + | + | + | + | + |
| SBGNview [33] | V | + | - | - | + | - | - | + |
| SBGNViz [125] | V | + | - | + | - | - | - | + |
| SBML Layout Viewer [29] | V | + | + | + | - | - | - | + |
| VISIBIOweb<sup>a</sup> [32] | V | + | - | - | - | - | - | + |
| yEd [158] + ySBGN [8] | E | + | - | - | - | - | - | - |

<sup>a</sup>Not actively maintained

## IV. SBGN-ED

### A) SBGN-ED overview

SBGN-ED [26, 65] is implemented as an Add-on (extension) for Vanted [67, 121]. Vanted is a software for visualising and analysing biological networks and associated experimental data. The software enables the import, creation, and editing of biological networks, the integration and visualisation of experimental data sets in the context of underlying networks as well as the visual exploration of networks and data. It also offers various possibilities for statistical data analysis and simulation. Visualisations of networks and data can be extensively customised by the user. The described functions are provided by Vanted itself or through corresponding Addons, and several Add-ons for Vanted exist such as for centrality analysis in networks (CentiLib [57]) or for working with SBGN (SBGN-ED).

Figure 2 shows SBGN-ED tabs when SBGN-ED is integrated in Vanted. There is a tab for each of the three SBGN languages and for the SBGN Bricks, a Tools tab, and an Examples tab, which provides example maps for all three languages. The tabs for each of the three SBGN languages contain interaction elements for all SBGN glyphs described in the specifications, which can be used to create and edit SBGN maps. The Bricks tab contains interaction elements for defined SBGN bricks which represent common building blocks for biological processes, following the idea described in [66, 124]. These bricks ease the assembly of SBGN maps and can be used to create SBGN maps when specific biological processes are needed on the map. If necessary, additional SBGN glyphs from the three tabs for the SBGN languages can be added to complete a map.

**Fig. 2:**
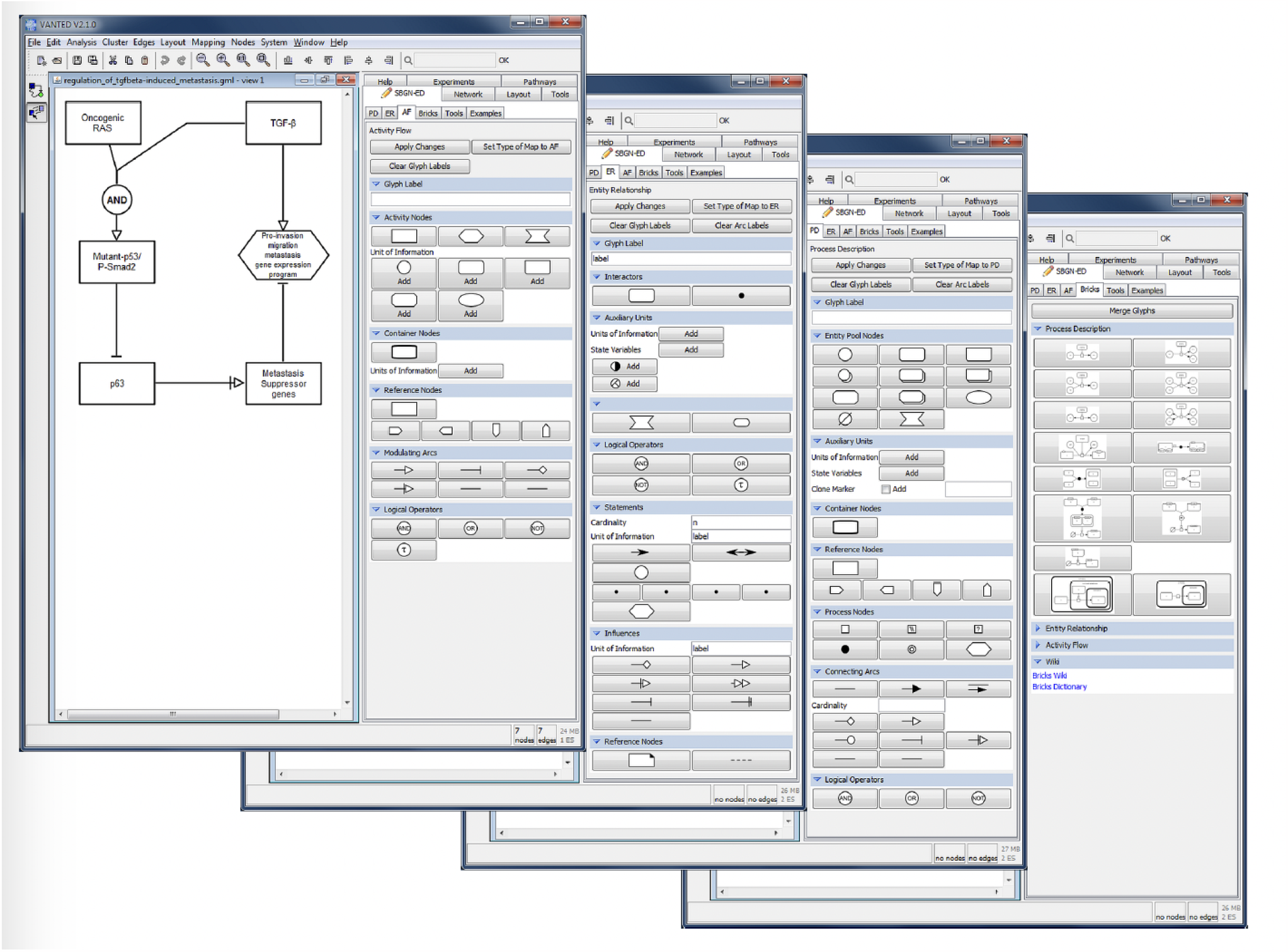
Screenshots of SBGN-ED integrated as an Add-on in Vanted. Left: Tab for the AF language and an AF map showing the regulation of TGF*β*-induced metastasis; Centre left: Tab for the ER language; Centre right: Tab for the PD language; Right: Tab for the SBGN Bricks.

SBGN maps which have been created or edited can be validated according to the SBGN specifications, i. e., are the glyphs on the map presented following the rules of the according specification, and are the glyphs connected by the correct arcs. The validation is provided in the Tools tab.

In addition to creating, editing, and validating SBGN maps, SBGN-ED also allows translating maps (or diagrams). This functionality can also be found on the Tools tab. SBGNED can translate KEGG diagrams downloaded with or shown in Vanted from maps in KEGG style (KEGG pathway database [69, 70]) into maps in SBGN PD style including automatic layout of the resulting SBGN map based on the initial KEGG diagram layout [27]. Furthermore, SBGN-ED can translate SBML models opened and visualised in Vanted into SBGN PD maps. And finally, SBGN-ED provides a function to translate SBGN maps from the Process Description language into the Activity Flow language [152].

### B) Building aesthetically pleasing SBGN visualisations

We introduce a five-step process for crafting aesthetically pleasing SBGN visualisations utilising SBGN-ED. This structured approach ensures that your maps are accurate and visually engaging. Those steps are also shown in Fig. 3 and 4.

**Fig. 3:**
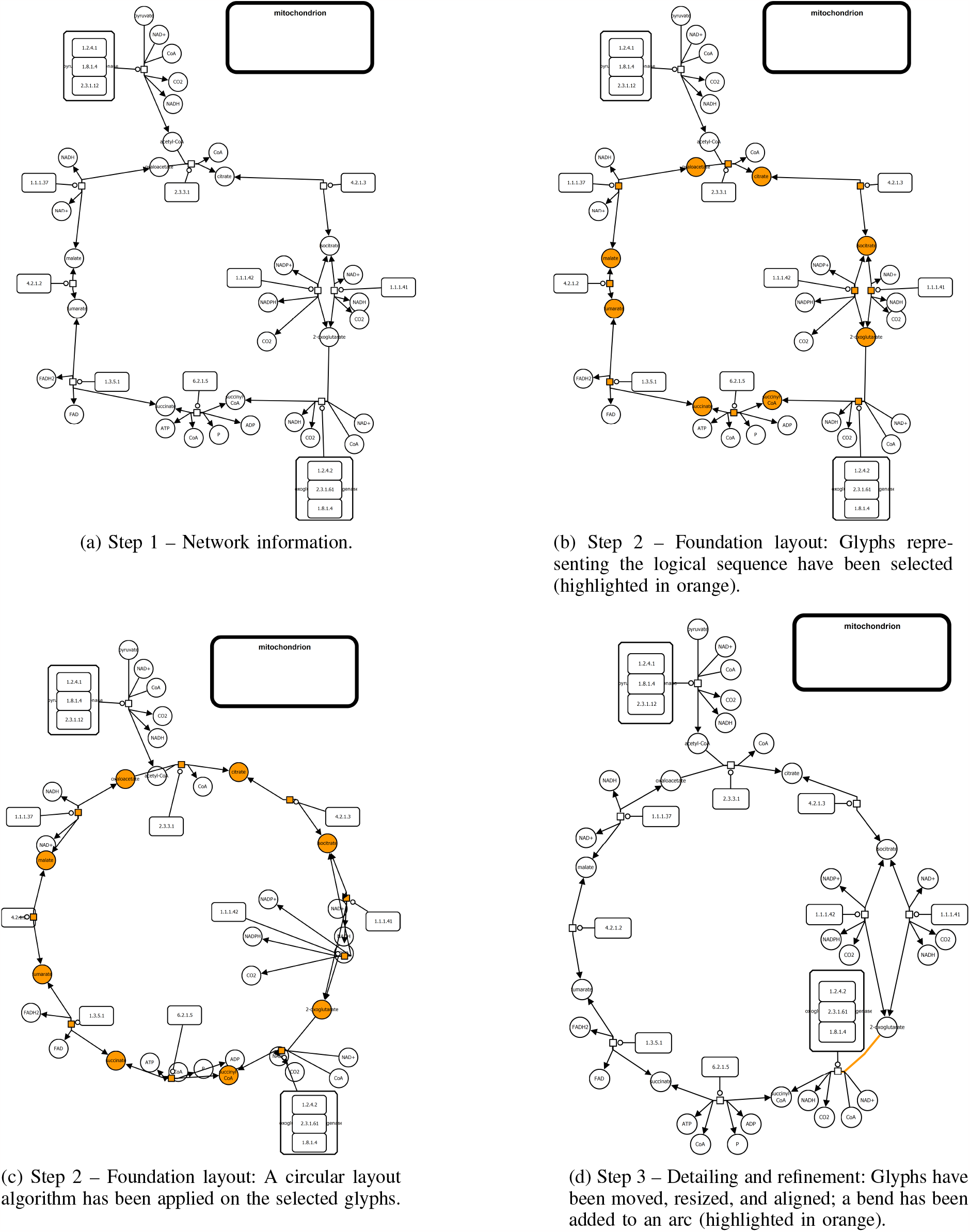
Steps 1 to 3 of the five-step process for building aesthetically pleasing SBGN visualisations utilising SBGN-ED (see Fig. 4 for the last steps). The steps show the creation of a visualisation for the TCA cycle from the MetaCrop [56, 133] database.

**Fig. 4:**
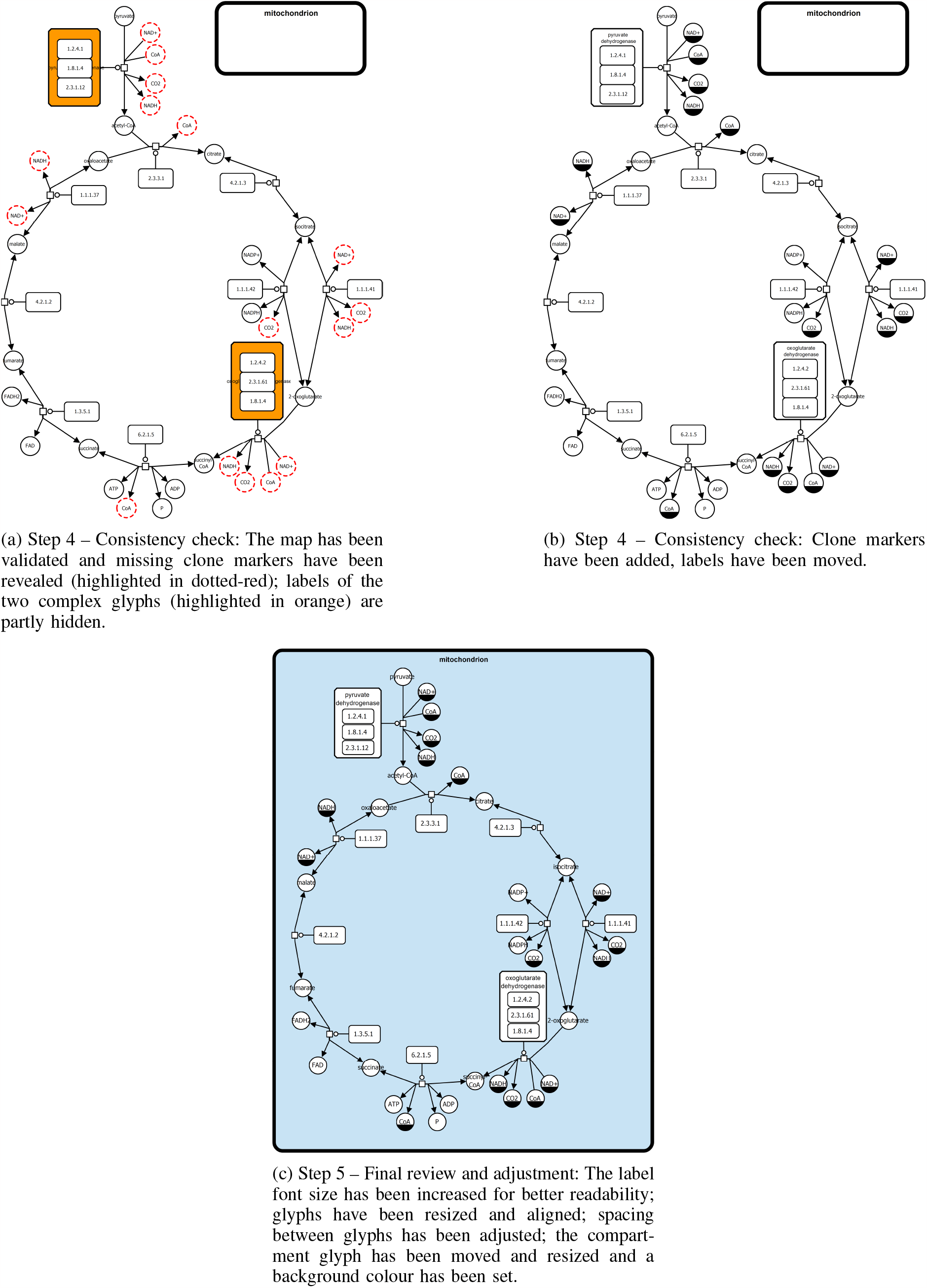
Steps 4 to 5 of the five-step process for building aesthetically pleasing SBGN visualisations utilising SBGN-ED (see Fig. 3 for the first steps). The steps show the creation of a visualisation for the TCA cycle from the MetaCrop [56, 133] database.

1. *Network information:* Begin by including the elements of your SBGN map in SBGN-ED (see Fig. 3a). Choose the right type of map (PD, ER or AF), and either manually build the network using the editor function of Vanted or the SBGN bricks in the Bricks tab, or load a network from an SBGN-ML [14, 150] file, or translate it from a SBML [62, 73] file or a KEGG [69, 70] file.
2. *Foundation layout:* Begin by sketching a basic layout of your SBGN map in SBGN-ED (see Fig. 3b and 3c). This involves placing the primary nodes and pathways in a logical sequence to reflect the biological process accurately. Typical layout algorithms can be used to support the creation of the basic layout, and parts of the network (map) can be drawn differently by marking the relevant set of nodes (glyphs) and edges (arcs) and apply the layout only to those nodes.
3. *Detailing and refinement:* Move elements or groups of elements to their final position (see Fig. 3d). Bends can be added or removed from edges, compartment nodes can be resized, and elements can be aligned by using functions of Vanted. During this phase, focus on refining the placement of these elements to enhance clarity and flow.
4. *Consistency check:* Review your diagram for consistency in symbols, labels, and layout (see Fig. 4a and 4b). Consistency is key in SBGN maps to ensure that they are easily interpretable and meet the standard notation guidelines. SBGN-ED provides syntax checking which highlights errors in the map such as reoccurring species without clone markers, forbidden connections between elements and more. Also review the meaning of your SBGN map.
5. *Final review and adjustment:* Conduct a final review of your map (see Fig. 4c). Make adjustments for aesthetic balance and readability. This might include tweaking the spacing, alignment, and overall layout to ensure that your visualisation is both informative and visually appealing.

By following these steps in SBGN-ED, it is possible to create SBGN visualisations that are not only scientifically rigorous but also aesthetically pleasing, enhancing both understanding and engagement with the audience.

### C. Additional features of SBGN-ED

SBGN-ED leverages the capabilities of the Vanted system to enhance the presentation and analysis of biological networks. One notable feature is its ability to export SBGN maps as clickable HTML images. This functionality facilitates building databases with interactive exploration of biological pathways, allowing users to click on different elements of the map to access more detailed information or related data. Examples are the MetaCrop database (see https://metacrop.ipkgatersleben.de/) and QSDB, the Quorum Sensing DataBase (see http://qsdb.org), where the clickable images for the graphical user interface have been built with SBGN-ED and link to information from additional databases.

SBGN-ED also supports the mapping of experimental data onto the elements of a SBGN map [36]. This feature is particularly useful, as it allows to overlay data directly onto the graphical representation of biological processes. It thereby provides a more comprehensive and integrative view of how experimental observations align with the theoretical framework of biological networks, which can enhance the understanding of biological systems. Figure 5 shows an example.

**Fig. 5:**
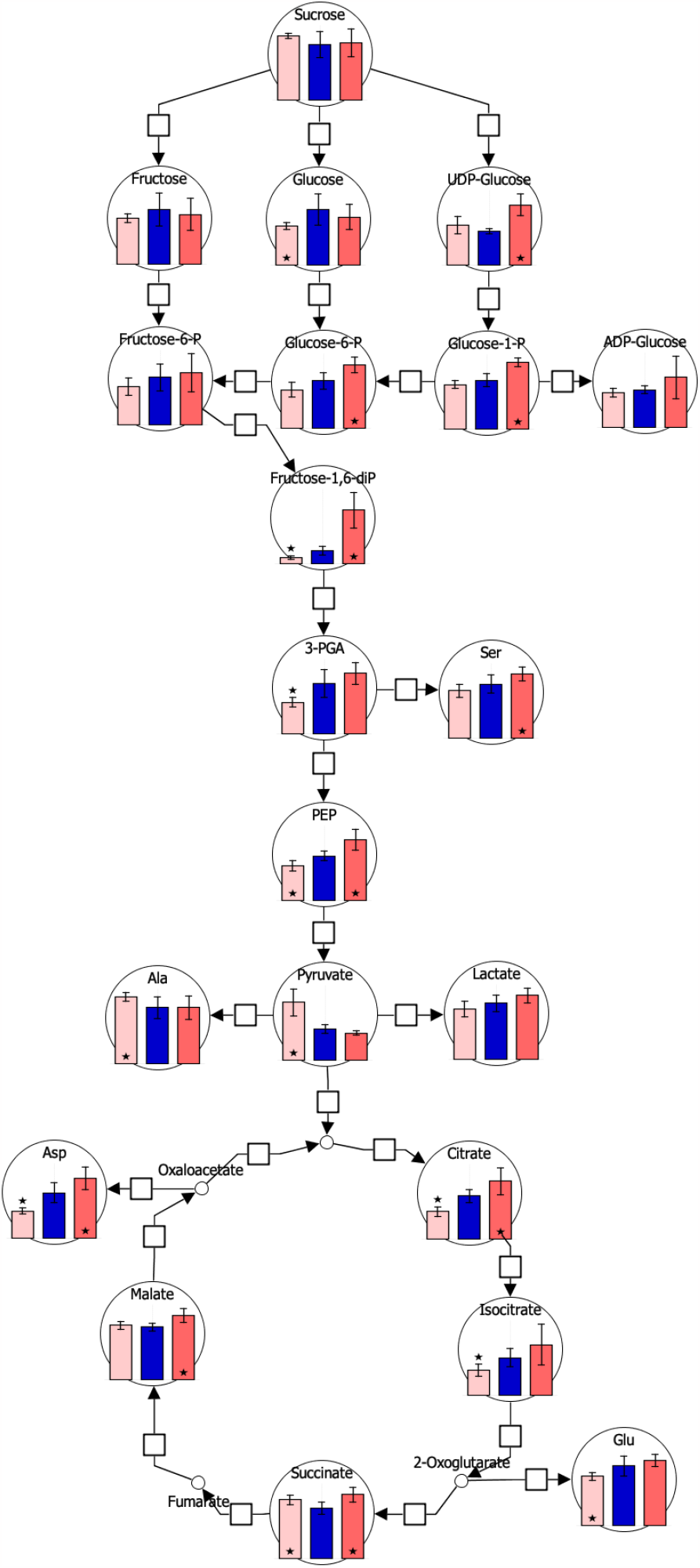
SBGN map of glycolysis and citric acid cycle with metabolite data from an experiment.

The exploration of large networks (examples include whole cell [155] or GSMM modelling [160, 161]) is supported by another Vanted Add-on, LMME [1]. LMME supports a process that begins by breaking down a large graph into multiple subgraphs, then identifying and visualising the connections between them. This initial visualisation gives a broad perspective; from there, more detailed subviews can be created to examine the subgraphs and their interactions more closely. These decompositions might be preset or calculated using existing or custom-developed techniques.

SBGN-ED has also been included into pipelines to provide SBGN map visualisation, examples are Path2Models [20] and the pipeline to create and analyse the COVID-19 Disease Map [108, 109].

## V. Conclusion

Emphasising the need for precision, engagement, information clarity, and aesthetic appeal in graphical data representations, we underscored the significance of the Systems Biology Graphical Notation (SBGN) and of proper layout of SBGN maps. As a standardised approach, SBGN effectively illustrates processes and networks in systems biology. We investigated the intricacies of SBGN, focusing on the defined graphical elements and the essential rules for connecting these elements to represent knowledge effectively. While the structure of the map elements is predetermined, their placement is crucial for creating visualisations that are not only useful but also visually appealing. Our presentation encompassed essential layout considerations and guidelines specific to SBGN maps. Furthermore, we introduced workflows for utilising SBGN-ED, a tool instrumental in designing visually engaging and exploratory SBGN visualisations. This comprehensive overview and the provided guidelines are aimed at enhancing the quality and impact of visual representations in the field of systems biology.

In this paper we focused on 2D visualisations. With the rise of immersive environments (such as head-mounted displays, fishtank VR, holographic displays, etc.) and immersive analytics as a new research field [23, 82, 83, 93], visualisations may move further into 3D [76] or transition smoothly between 2D and 3D (transitional hybrid interfaces) [2, 142, 159]. There have been early approaches for general biological networks in 3D, for example, using CAVEs [110, 119, 122, 141], approaches stacking biological networks in 2.5D (and thereby moving towards 3D) [18, 19, 44], as well as the before mentioned transitional interfaces. However, good, ideally mental-map preserving interactive 3D layouts of SBGN maps are still an open research question.

## Notes

### Competing Interest Statement

The authors have declared no competing interest.

